# Metformin improves cognition of aged mice by promoting cerebral angiogenesis and neurogenesis

**DOI:** 10.1101/2020.03.25.006767

**Authors:** Xiaoqi Zhu, Junyan Shen, Shengyu Feng, Ce Huang, Zhongmin Liu, Yi. Eve. Sun, Hailiang Liu

## Abstract

The cerebral microvasculature is essential for preservation of normal cerebral function. Age-related decreases of neurogenesis and cognitive function are accompanied by reduced blood flow and a decline in neural stem cell (NSC) number. Here, we report that metformin administered by tail vein injection enhanced cognition in aged but not young mice in a dose-dependent manner. Further, metformin restored cerebral blood flow and brain vascular density and promoted neurogenic potential of the subependymal zone/subventricular zone both in vivo and in vitro. RNA-Seq result indicated that metformin could enhance glycolysis in blood, with an increase in relative mRNA expression of the enzyme in the glycolysis pathway from hippocampal tissue of metformin-treated mice. Mechanistically, glyceraldehyde-3-phosphate dehydrogenase (GAPDH), a key enzyme in the glycolysis pathway, may contribute to angiogenic and neurogenic potentials of NSCs. Interestingly, examination of peripheral blood mononuclear cells from people of various ages showed that mRNA expression of GAPDH gradually decreased with age, while its expression level positively correlated with cognitive levels. Our results indicate that metformin represents a candidate pharmacological approach for recruitment of NSCs in aged mouse brain by enhancing glycolysis and promoting neurovascular generation, a strategy that might be of therapeutic value for anti-aging in humans.

**Graphical Abstract:** 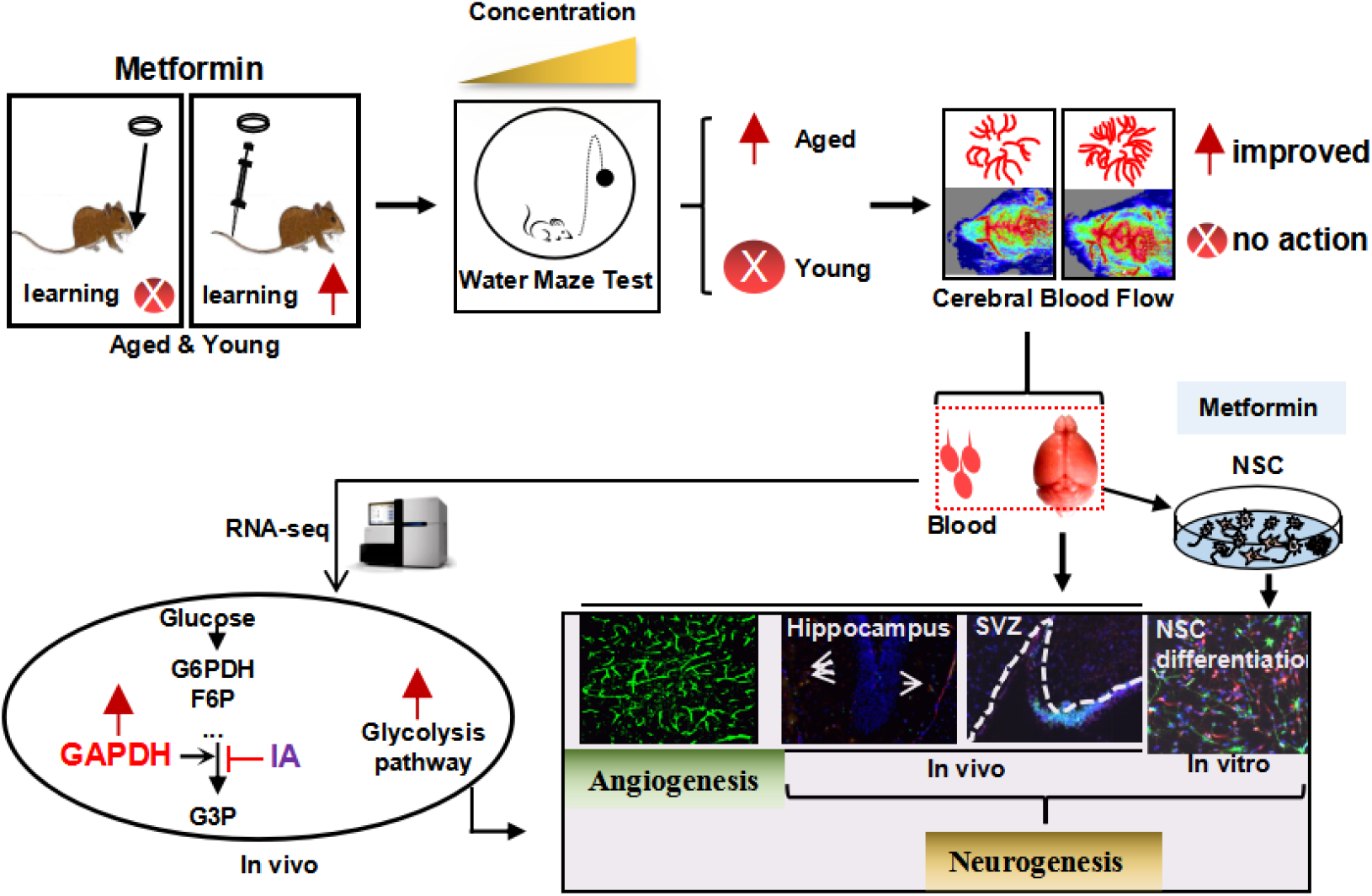

## Introduction

The brain is the most metabolically active organ in the human body. It only accounts for 2% of body mass, but consumes 20%–25% of the body’s total energy requirements. The cerebromicrovascular system is crucial for brain health and function, and constantly supplies nutrients and oxygen as well as effective washout of metabolic waste products [1]. However, compelling evidence obtained from both elderly patients and rodent models shows that aging significantly impairs neurovascular coupling responses [2–4]. Importantly, age-related neurovascular impairments have been linked to impaired cognitive function and gait abnormalities.

Increasing emerging evidence indicates that glucose metabolism controls endothelial cell proliferation, migration, and neovascularization generation [2, 5]. Additionally, blood vessel function is required for efficient neural stem cell (NSC) proliferation and differentiation, providing oxygen or a local source of signaling molecules secreted from endothelial cells and also delivering systemic regulatory factors that may regulate NSC metabolism [6]. Thus, therapeutic interventions that restore neurovascular function in elderly patients have the potential to improve a range of age-related neurological deficits. Metformin, a drug approved to treat diabetes, appears to target many aging-related mechanisms, such as inhibition of mitochondrial complex 1 in the electron transport chain, and consequent reduction of endogenous production of reactive oxygen species (ROS) [7]. In addition, metformin was shown to stimulate glycolytic lactate production in cultured primary rat astrocytes [8]. Unexpectedly, metformin can also recruit NSCs to enhance neural function and restore central nervous system remyelination capacity by rejuvenating aged stem cells [9, 10]. These findings arouse our interest to determine whether age-related declines of cognitive level, vascular integrity, and neurogenic niche can be restored by administering metformin. Overall, our results suggest that metformin has beneficial effects by tail vein injection (not oral administration) on cognitive levels in aged mice. These effects are associated with restoration of vascular integrity, producing a richer cerebral blood flow as well as activation of neurogenesis in the subependymal zone/subventricular zone (SEZ/SVZ). Furthermore, mRNA expression levels of glyceraldehyde-3-phosphate dehydrogenase (GAPDH), a key glycolysis enzyme, gradually decreased with age, and positively correlated with cognitive levels. Meanwhile, metformin administration enhanced glycolysis through increased mRNA expression of GAPDH, which ultimately increased angiogenesis and neurogenic potential of NSCs. Our results indicate that metformin represents a candidate pharmacological approach for recruitment of NSCs in aged mouse brain. Further, metformin restores neurovascular integrity by enhancing glycolysis, a strategy that might be of therapeutic value for anti-aging in humans.

## Results

### Metformin by tail vein injection improved cognitive levels of aged mice

As aging is frequently accompanied by cognitive impairments, we examined the effect of metformin administration by tail vein injection on spatial learning and memory of aged mice. Mice aged 4, 10–12, or 20 months were treated with metformin by tail vein injection every 2 days for 1 month. Subsequently, Morris water maze (MWM) testing was used to examine the effect of metformin on spatial learning and memory. Five-day acquisition trials were then conducted using a hidden platform test. Short-term memory retention was tested 24 h after 5 days of training. Decreased escape latency was detected among 10–12 month and 20-month metformin-treated mice (Figure 1A and 1D). In this probe test, the metformin-treated group showed a significantly enhanced ability to reach the virtual platform, as measured by first time-to-platform (Figure 1B and 1E). Both control and metformin groups exhibited similar swim speeds to the virtual platform (Figure 1C and 1F), suggesting comparable vision and motivation between the two groups. To distinguish the effect of different metformin doses on cognitive function, 20-month mice were treated with metformin at different concentrations using the same procedure described above. The results showed that the beneficial effect of metformin on decreased escape latency of aged mice exhibited a dose-dependent tendency (Figure 1D). In the probe test, first time-to-platform among the different metformin doses also exhibited a dose-dependent tendency (Figure 1E). However, 4-month-old mice treated with metformin did not exhibit improved cognitive function, as escape latency in the metformin-treated group was not different than the control group during the five-day training period (Supplementary Figure 1A). In the probe test, first time-to-platform was not significantly decreased in the metformin-treated group (Supplementary Figure 1B). Both the control and metformin groups exhibited similar swim speeds to the virtual platform (Supplementary Figure 1C), suggesting that metformin did not improve the cognitive function of young mice. Collectively, these results indicate that metformin can enhance spatial learning and memory function of aged mice, but not young mice, and may exhibit a dose-dependent tendency.

**Figure 1.**
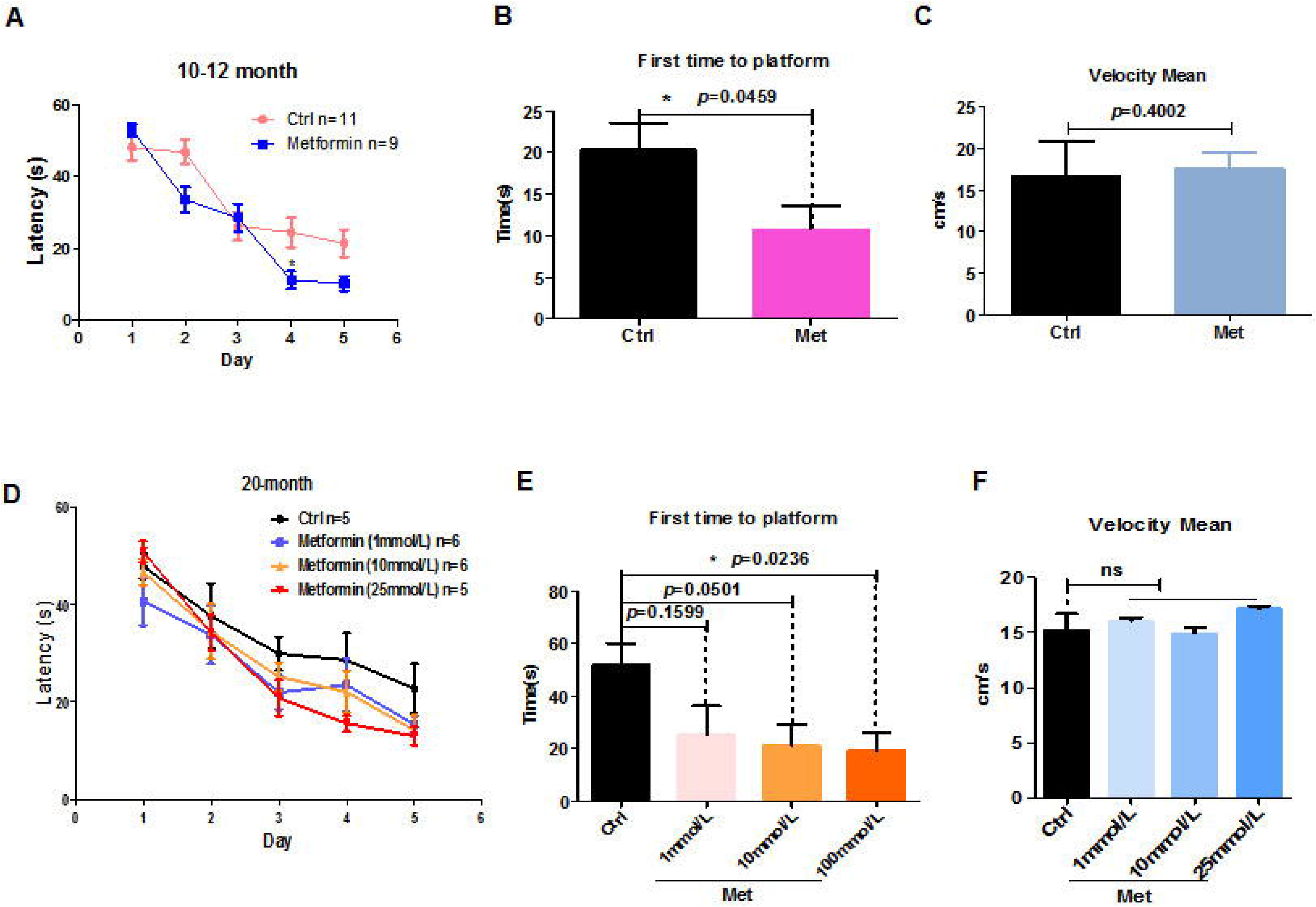
Metformin improved spatial learning and memory of older mice in the Morris water maze test. (A) Escape latency of 10–12-month mice treated with metformin (*n* = 9) was significantly shorter compared with the control group (*n* = 11). (B) Probe tests conducted 24 h after the acquisition phase indicated that the first time-to-platform of 10–12-month-old mice treated with metformin was shorter than the control group. (C) No difference in swim speed was observed between control and metformin-treated groups. (D) Mean escape latency during 5 training days in 20-month-old mice treated with different doses of metformin. (E) Probe tests conducted after 24 h of the acquisition phase indicated that first time-to-platform was shorter in 20-month-old mice treated with higher doses of metformin than the control group. (F) No differences in swim speed were observed between control and metformin-treated groups (number of mice in control group, *n* = 5; in 1 mMol/L group, *n* = 6; in 10 mMol/L group, *n* = 6; in 25 mMol/L group, *n* = 5; Ctrl: Control; Met: Metformin). The overall significance between two groups was determined by Student’s *t*-test. * *p* < 0.05, ** *p* < 0.01, *** *p* < 0.001, ns, not significant.

### Metformin restores brain vascular integrity in aged mice

Aging significantly impairs neurovascular coupling responses. Such age-related neurovascular deterioration is accompanied with a sharp decrease in cerebral blood flow (CBF) [11]. To determine whether the beneficial effect of metformin on spatial learning and memory ability was associated with increased CBF, Doppler laser blood stream detector was used to examine CBF in mice treated with metformin. The results indicated that fold-change relative to the control group in CBF responses to contralateral whisker stimulation was significantly increased in 10–12 month- and 20-month-old aged mice (Figure 2A and 2B). Moreover, CBF of 20-month-old mice treated with different dose of metformin exhibited a dose-dependent tendency (Figure 2B). However, CBF of 4-month-old mice treated with metformin was not significantly increased compared with the control group (Supplementary Figure 1D). To further examine the effects of metformin on cerebral microvasculature, immunofluorescence staining was performed. Our results showed that the number of branchpoints in the hippocampus and cerebral cortex (labeled green) were significantly increased in the metformin-treated group (Figure 2C). These results demonstrate that metformin can restore cerebromicrovascular integrity impairments and improve cognitive function in aged mice.

**Figure 2.**
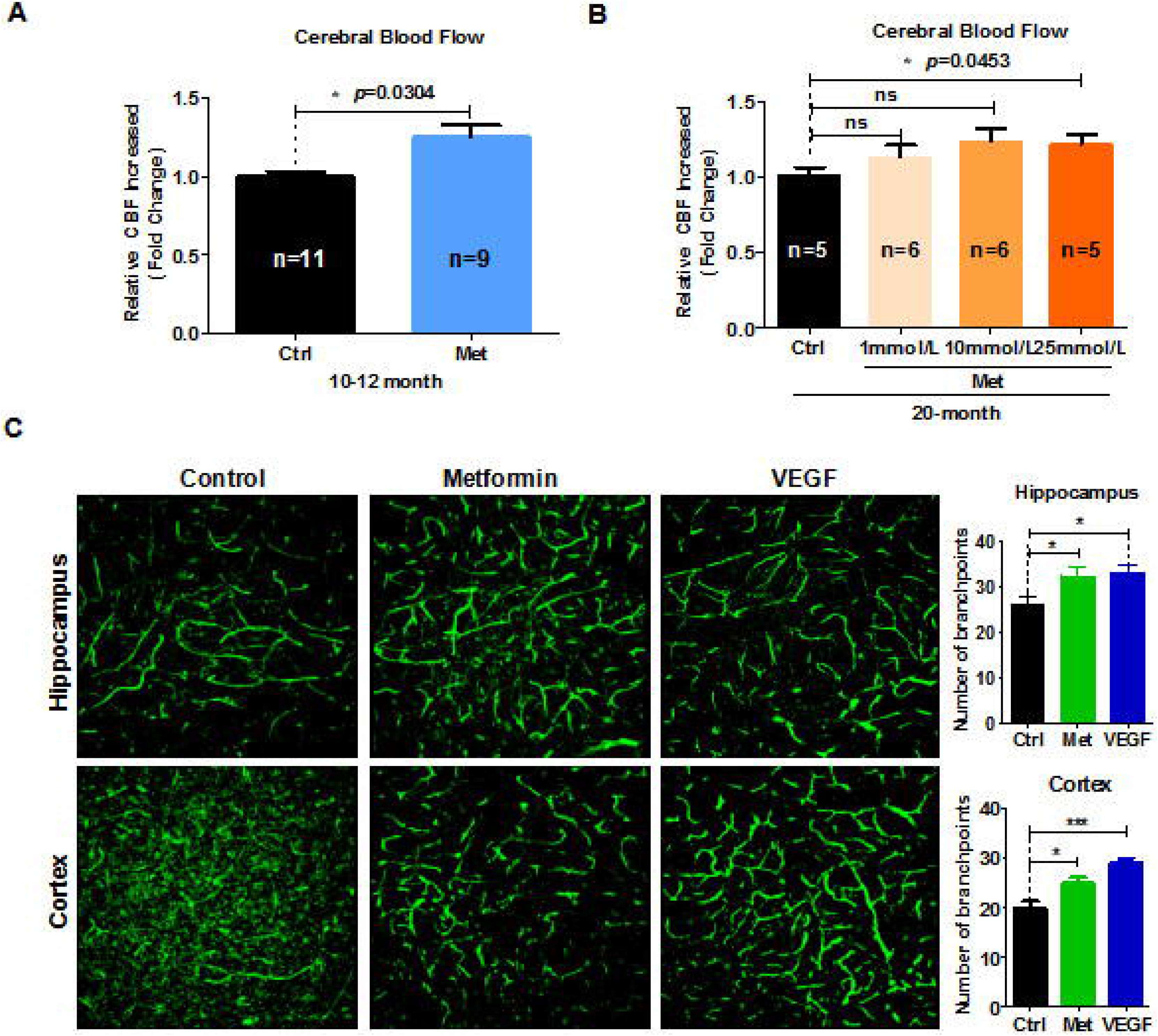
Metformin restores brain vascular integrity of old mice. (A) Fold-change relative to the control group (*n* = 11) in cerebral blood flow (CBF) response to contralateral whisker stimulation was significantly increased in 10–12-month-old mice treated with metformin (*n* = 9). (B) Fold-change relative to the control group in CBF response to contralateral whisker stimulation was significantly increased in 20-month-old mice treated with metformin, and exhibited a dose-dependent tendency (number of mice in control group, *n* = 5; in 1 mMol/L group, *n* = 6; in 10 mMol/L group, *n* = 6; in 25 mMol/L group, *n* = 5). (C) (left) 3D reconstruction of the hippocampus dentate gyrus and cortex zone vasculature generated by confocal imaging of 100 μm thick sections. (right) Quantification of blood vessel branchpoint per field in the hippocampus dentate gyrus and cortex of metformin, vascular endothelial growth factor (VEGF), and control groups (every treated group *n* ≥ 3). Scale bar = 100 μm. Ctrl: Control; Met: Metformin. The overall significance between two groups was determined by Student’s *t*-test. * *p* < 0.05, ** *p* < 0.01, *** *p* < 0.001, ns, not significant.

### Metformin rejuvenation of neurogenic potential in aging mice

The vasculature can influence NSC proliferation and differentiation, as it provides a local source of signaling molecules secreted from endothelial cells or oxygen as well as delivers systemic regulatory factors and possibly regulates NSC metabolism [12]. Considering that aging causes reduced numbers of progenitor cells, and that metformin can rejuvenate brain vascular integrity of aged mice, we next examined neurogenic potential (e.g., SRY-box 2 [Sox2]^+^ stem cells, doublecortin [DCX]^+^ newborn neurons) of aged mice by analyzing sagittal SVZ sections of metformin-treated brain. Our results show that Sox2^+^ NSC numbers were significantly increased in the SVZ (Figure 3A and 3C). In addition, DCX^+^ cell numbers were also increased in SVZ areas of the metformin-treated group (Figure 3A and 3B). To determine whether the increase in neural stem and progenitor cells could produce a subsequent change in SVZ neurogenesis in the metformin-treated group, aged mice were injected with bromodeoxyuridine (BrdU) three-times after metformin treatment at 24-h intervals to label actively dividing cells. The mice were analyzed for BrdU^+^/NeuN^+^ cells to quantify newborn neurons. Our results showed that metformin increased the number of BrdU^+^/NeuN^+^ cells in the SVZ (Figure 3D and 3E). To further validate these results, we microdissected the lateral ventricle walls (ependymal zone together with SEZ/SVZ) covering the striatum, and dissociated the tissue into a single cell suspension to obtain NSCs. To confirm that metformin treatment promoted neurogenesis of NSCs, we examined spontaneous differentiation of primary NSCs treated with metformin. First, a single-cell suspension of NSCs was plated onto laminin-coated coverslips at a density of 2 × 10^5^ cells/cm^2^. When cell confluence reached approximately 80%, basic fibroblast growth factor (bFGF) was withdrawn and NSCs were treated with metformin. After 7 days of continuous treatment with metformin, cells were fixed with 4% paraformaldehyde (PFA) and immunostained with glial fibrillary acidic protein (GFAP) and class III beta-tubulin (Tuj1) antibodies. The results showed that the ratio of Tuj1^+^ cells to total cells in the metformin-treated group was significantly increased (Figure 3F and 3H), while the ratio of GFAP^+^ cells to total cells was slightly decreased (Figure 3F and 3G). These results suggest that upon differentiation, metformin-treated NSCs differentiated into more neurons compared with control NSCs, indicating increased neurogenic potential. Taken together, these observations suggest that metformin promotes the neurogenic potential of NSCs in the SVZ zone both *in vivo* and *in vitro*.

**Figure 3.**
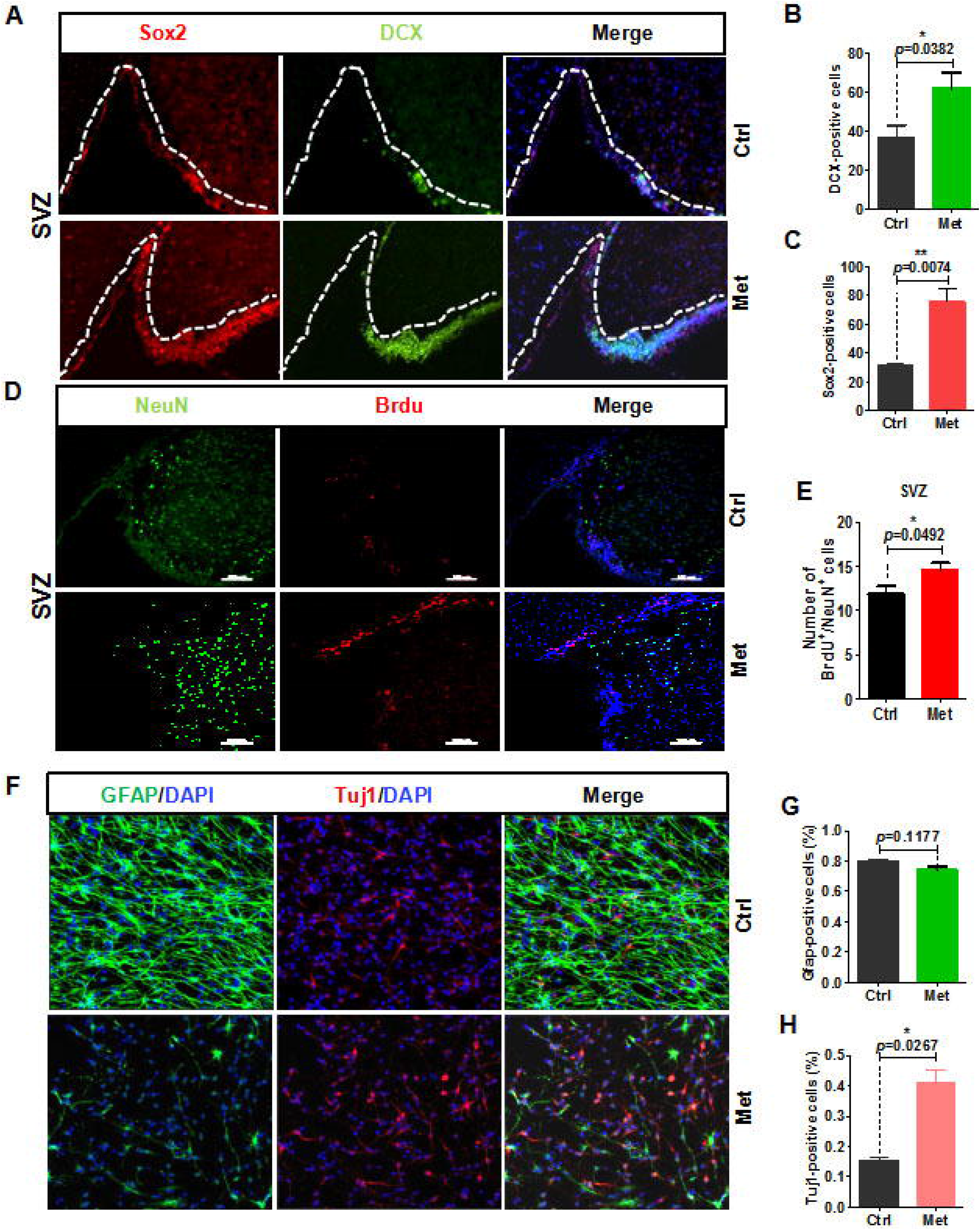
Metformin rejuvenation of neurogenic potential in aged mice. (A) The subventricular zone (SVZ) of old mice treated with metformin (*n* ≥ 3) immunostained with the neural stem cell marker, Sox2, and newborn neuronal marker, GFAP. Right: Quantification of (B) Sox2^+^ cells and (C) DCX^+^ cells. (D) SVZ of old mice immunostained with the neuronal cell marker, NeuN, and cell proliferation marker, BrdU. Right: Quantification of (E) BrdU^+^/NeuN^+^ cells to quantify newborn neurons. (F) Representative fields of GFAP and Tuj1 immunofluorescence staining of cultured neural stem cells treated with metformin after 7 days of spontaneous differentiation. Right: Statistical analysis of percentages of (G) GFAP^+^ cells and (H) Tuj1^+^ cells. Scale bar = 100 μm. Ctrl: Control; Met: Metformin. The overall significance between two groups was determined by Student’s *t*-test. * *p* < 0.05, ** *p* < 0.01, *** *p* < 0.001, ns, not significant.

### Metformin enhanced glycolysis in blood of aged mice

To examine the molecular mechanisms underlying the beneficial effects of metformin on cognitive function of aged mice, we performed whole genome transcriptomic analyses combined with weighted gene co-expression network analysis (WGCNA) of peripheral blood from mice treated with metformin (Figure 4A). Differential gene expression clearly demonstrated that compared with the control group, glycolysis/gluconeogenesis, Forkhead box (FOXO) signaling, and 5’ AMP-activated protein kinase (AMPK) signaling pathways were upregulated in the metformin-treated group (Figure 4B and 4C). In addition, we found decreased plasma fructosamine in metformin-treated mice (Supplementary Figure 2). Increasing emerging evidence indicates that glucose metabolism controls endothelial cell proliferation, migration, and neovascularization generation. Thus, we examined relative mRNA expression of enzymes in the glycolysis pathway of metformin-treated hippocampus tissue by quantitative real-time polymerase chain reaction (qRT-PCR). Our results showed that relative GAPDH and platelet isoform of phosphofructokinase (PFKP) mRNA expression were upregulated (Figure 4D and 4E).

**Figure 4.**
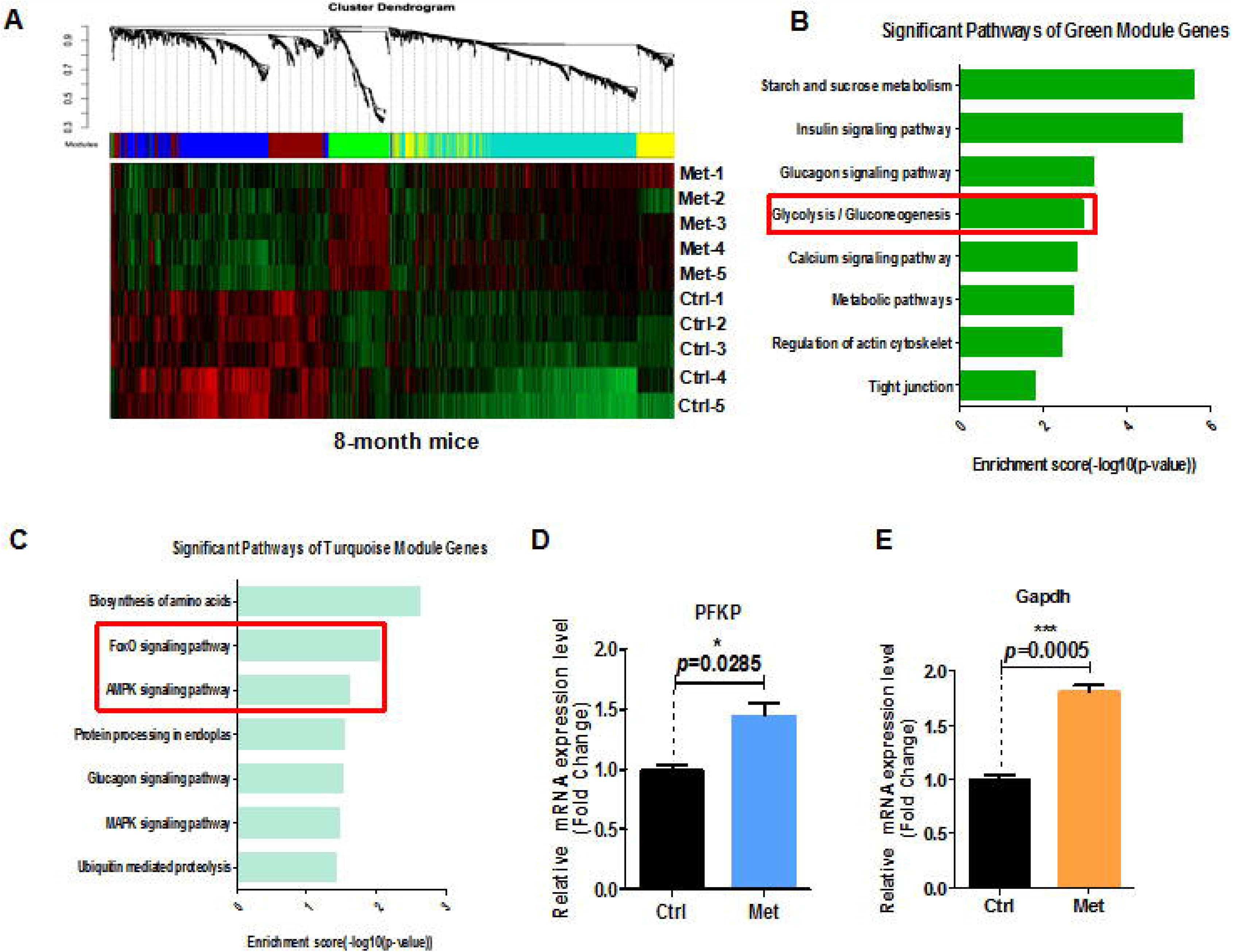
Transcriptome analysis of blood of 8-month-old mice treated with metformin. (A) Heatmap of differentially expressed genes in the blood of 8-month-old mice treated with metformin. (B) and (C) KEGG pathway analysis of green and turquoise modules. (D) Relative PFKP mRNA expression in hippocampal tissue from metformin-treated mice. (E) Relative GAPDH mRNA expression in hippocampal tissue from metformin-treated mice (every treated group *n* ≥ 3). Ctrl: Control; Met: Metformin. The overall significance between two groups was determined by Student’s *t*-test. * *p* < 0.05, ** *p* < 0.01, *** *p* < 0.001, ns, not significant.

### GAPDH was the key enzyme in the glycolysis pathway positively associated with increased cognitive levels in humans

Age-related decreases in brain glucose uptake exceed that of oxygen use, resulting in loss of brain aerobic glycolysis and blood flow. RNA-Seq results indicated that metformin could enhance levels of glycolysis of aged mice both *in vivo*. GAPDH is an important enzyme for glycolysis that converts glyceraldehyde-3-phosphate derived from glucose to 1, 3-bisphosphoglycerate in the presence of NAD^+^ [13]. Apart from this basic function, GAPDH has been suggested to play multifunctional roles in biological processes including apoptosis [14], endocytosis [15], DNA replication [16], and DNA repair [17]. Research has shown that GAPDH, the enzyme separating lower and upper glycolysis, is the rate-limiting step in the glycolysis pathway. Further, levels of fructose (1, 6) bisphosphate are predictive of the rate and control points of glycolysis [18]. Thus, we speculated that GAPDH is the key enzyme in the glycolysis pathway enhanced by metformin. To test this hypothesis, NSCs were treated with metformin and iodoacetic acid (IA), an inhibitor of GAPDH [18]. Relative GAPDH mRNA expression was significantly upregulated in NSCs from metformin-treated groups, while IA-treated groups showed significant downregulation (Figure 5A). Meanwhile, relative mRNA expression of the angiogenesis-related gene, vascular endothelial growth factor receptor 2 (VEGFR2), was also significantly upregulated in NSCs from metformin-treated groups, but significantly downregulated in IA-treated groups (Figure 5B). These results indicate that GAPDH was enhanced by metformin and served as a key factor to promote angiogenesis. To examine this effect of GAPDH in the context of neurogenesis, NSCs were similarly treated with metformin and IA. Relative mRNA expression of the neuronal marker, Tuj1, was significantly increased in the metformin-treated group and significantly downregulated in the IA-treated group (Figure 5C). Relative mRNA expression of the astrocyte marker, GFAP, was not significantly different in the metformin-treated group compared with the control group, but was significantly downregulated in the IA-treated group (Figure 5D). These results, which are consistent with those of spontaneous differentiation of metformin-treated primary neural progenitor cells, suggest that the key glycolytic enzyme, GAPDH, was enhanced by metformin, which promoted both angiogenesis and neurogenesis in NSCs. In addition, relative GAPDH mRNA expression levels in peripheral blood mononuclear cells from healthy individuals of young age (chronicle age: 20s and 30s) and old age (chronicle age: > 50s), as well as patients with Alzheimer’s disease, were gradually downregulated with aging (Supplementary Figure 3A and 3B). We next performed the Mini-Mental State Examination of old people (chronicle age > 70s) to determine correlation between cognitive function and GAPDH mRNA expression levels [19]. Our results indicate a positive association between age-dependent cognitive function and improvement in GAPDH levels (Supplementary Figure 3C).

**Figure 5.**
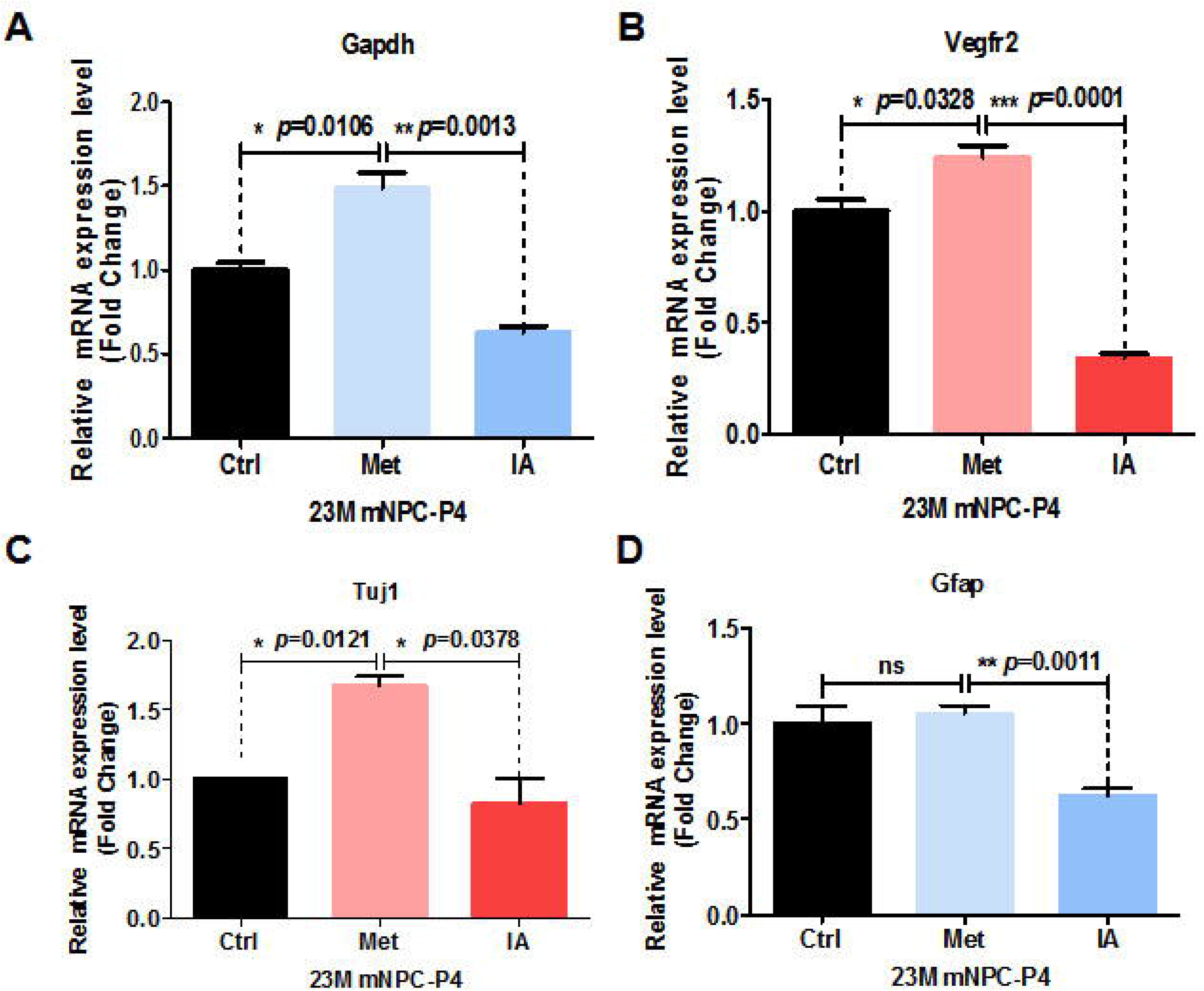
Metformin enhanced relative mRNA expression of GAPDH in the glycolysis pathway and was associated with relative mRNA-related angiogenesis and neurogenesis and increased expression. (A) Relative GAPDH mRNA expression of neural stem cells (NSCs) treated with metformin and iodoacetic acid (IA), as examined by qRT-PCR. (B) Relative VEGFR2 mRNA expression of NSCs treated with metformin and IA, as examined by qRT-PCR. (C) and (D), Relative Tuj1 (C) and GFAP (D) mRNA expression of NSCs treated with metformin and IA, as examined by qRT-PCR. Ctrl: Control; Met: Metformin. The overall significance between two groups was determined by Student’s *t*-test. * *p* < 0.05, ** *p* < 0.01, *** *p* < 0.001, ns, not significant.

### Treatment by tail vein injection results in better cognitive function than oral administration of metformin

Metformin is a drug approved to treat diabetes that appears to target many aging-related mechanisms. However, some studies have reported that long-term oral administration of metformin in old mice does not restore aging-related deficits [20]. To investigate the anti-aging effects of oral administration of metformin on old mice, 8-month-old mice were treated with metformin orally for 10 months, and the overall disability rate was recorded. The results showed that overall disability rate of metformin-treated mice was significantly lower than the disability rate of control mice (Figure 6A). This indicates that long-term use of metformin is associated with a marked impairment. To avoid this side effect of metformin by oral administration, we compared treatment of old mice by tail vein injection and oral administration. As aging is frequently accompanied by cognitive impairments, the MWM was used to examine the anti-aging effect of metformin treatment by different administration methods. The results showed that metformin treatment by tail vein injection to mice resulted in a better performance on first latency to platform compared with metformin by oral administration and the control group (Figure 6B). In the probe test, mice treated with metformin by tail vein injection showed a significantly enhanced ability to reach the virtual platform, as measured by first time-to-platform (Figure 6C). Both control and metformin groups exhibited similar swim speeds to the virtual platform (Figure 6D). Thus, we concluded that metformin treatment by tail vein injection is associated with better performance on spatial learning and memory tasks compared with oral administration of metformin.

**Figure 6.**
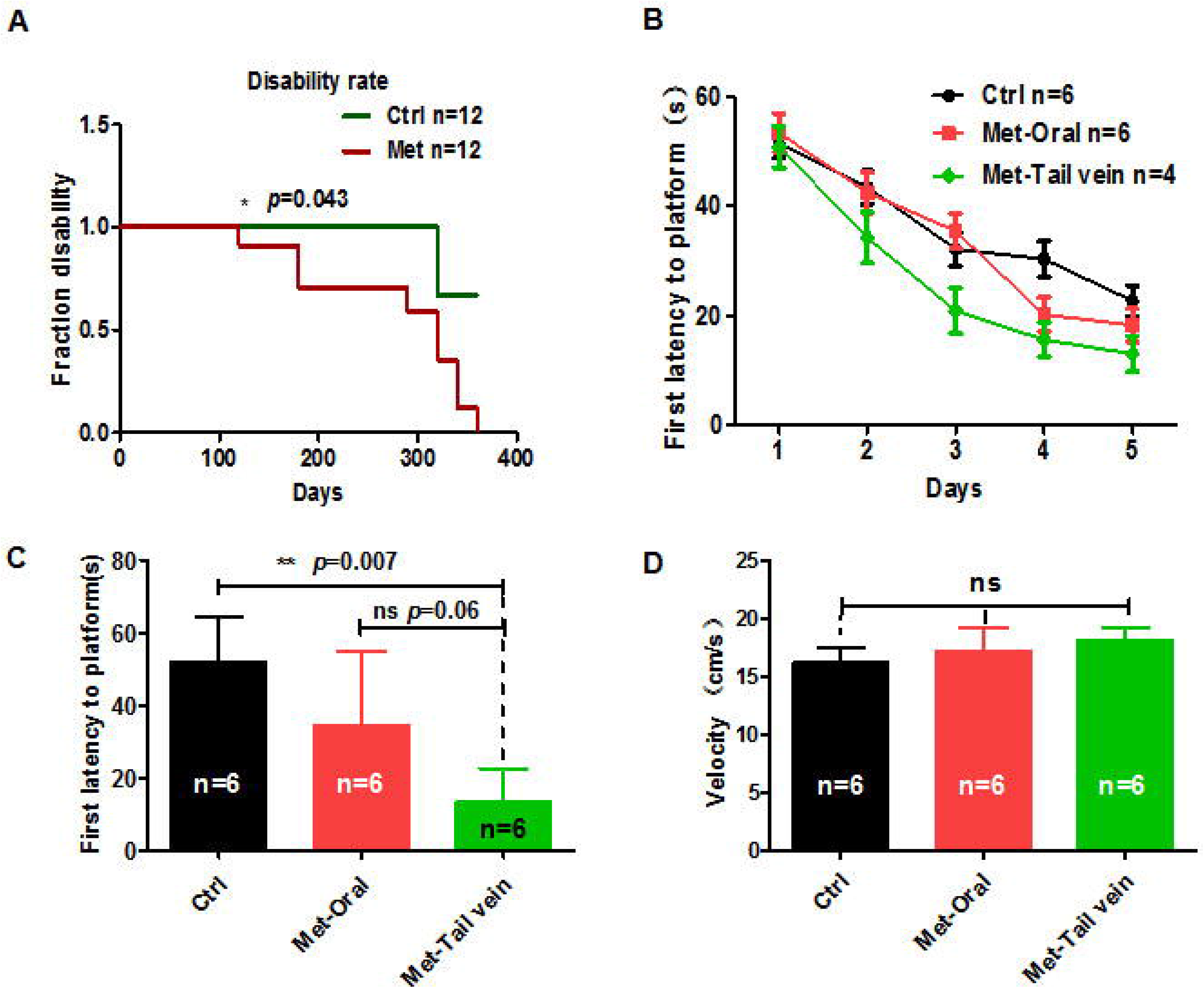
Mice treated with metformin by tail vein injection showed better performance in the Morris water maze test. (A) Mice treated with metformin (*n* = 12) for 10 months showed a considerable disability rate compared with the control group (*n* = 12). (B) Twenty-month-old mice treated with metformin by tail vein injection (*n* = 4) showed a better performance in escape latency than mice treated by oral administration (*n* = 6). (C) Probe tests conducted at 24 h after the acquisition phase indicated that the first time-to-platform of 20-month-old mice treated with metformin by tail vein injection was shorter than in mice treated by oral administration. (D) No difference in swim speed was observed between control and metformin-treated groups (tail vein injection or oral administration). Ctrl: Control; Met: Metformin. The overall significance between two groups was determined by Student’s *t*-test. * *p* < 0.05, ** *p* < 0.01, *** *p* < 0.001, ns, not significant.

## Discussion

Aging is a process involving slow deterioration of many homeostatic functions throughout an organism’s life. Middle age is a life period during which many physiological and psychological changes occur, leading to cognitive impairment and behavioral deterioration, and eventually deterioration of brain function [20–22]. Metformin, a drug approved to treat diabetes, appears to target many aging-related mechanisms. However, some studies hold a differing opinion, suggesting that oral administration of metformin cannot improve age-related impairments [23]. To determine whether metformin can rescue age-related impairments, we treated 8-month-old mice with metformin by oral administration for 10 months. Our results show that metformin significantly increased disability rate (Figure 6A), including cancer, cataract, and dermatitis, which is similar to most studies conducted on diabetics showing that long-term treatment with metformin causes a risk of vitamin B-12 deficiency and other side effects in patients with type 2 diabetes [24]. To avoid the side effects of metformin, we treated mice with metformin by tail vein injection, and used the MWM to examine the anti-aging effect of metformin treatment by different administration methods. Our results show that mice treated with metformin by tail vein injection have better performance on spatial learning and memory function with no other side effects. Thus, we treated mice with metformin by tail vein injection to examine its effect on cognitive function.

Studies have shown that metformin treatment via oral administration at young, middle, and advanced ages in male mice produces different results on healthy lifespan [25]. To investigate the varying effects of metformin on mice at different ages, we examined the effect of metformin treatment by tail vein injection of 4, 10–12, and 20 month-old female mice. Our results show that young mice treated with metformin exhibit no improvements in spatial learning and memory function. However, aged mice treated with metformin in similar procedures show significantly improved spatial learning and memory. These results are similar to research conducted by Rebecca M. Ruddy [26], showing that metformin could enhance neurogenesis in female mice. Thus, we concluded that these contradictory results of metformin treatment may be caused by gender or different administration methods of metformin treatment.

Glycolysis is a multi-step process that prepares the glucose molecule for oxidative phosphorylation and subsequent generation of energy. In the normal brain, glycolysis exceeds that required for the needs of oxidative phosphorylation. Because this process occurs in a setting with adequate oxygen available for oxidative phosphorylation, it is often referred to as aerobic glycolysis, and is a biomarker for metabolic functions broadly supporting biosynthesis and neuroprotection [27]. Numerous studies have shown that lactate production is beneficial for memory function during normal aging [28, 29]. Here, we demonstrated that metformin could enhance spatial learning and memory ability through glycolysis.

Glycolysis is a pivotal process during development and for adult vascular growth and development [30]. Thus, we performed immunohistochemical analyses and qRT-PCR to examine angiogenesis-related genes. Our results show that metformin significantly promoted angiogenesis, either through increased amounts of vascular endothelial cells or enhancement of angiogenesis-related gene expression. Many studies have shown that blood vessels are an essential component of stem cell niches, regulating the balance between precursor expansion and differentiation. Moreover, new blood vessels provide sufficient nutrients for NSCs, as co-culture of embryonic NSCs with immortalized endothelial cells *in vitro* increased NSC expansion and determined their fate toward neurons [31]. Other studies have shown that the pattern of angiogenesis in the cortex seems to be accompanied with initiation of neurogenesis *in vivo* [32]. Here, we demonstrated that metformin treatment significantly increased the number of Sox2^+^ and DCX^+^ NSCs and promoted neurogenesis. To determine whether aerobic glycolysis is indispensable for angiogenesis and neurogenesis in the brain, we treated cells with IA to inhibit GAPDH, the enzyme separating lower and upper glycolysis in the pentose phosphate pathway. Cells treated with IA exhibited significant suppression of genes associated with angiogenesis and neurogenesis, indicating that GAPDH is the rate-limiting step in the glycolysis pathway. Here, we demonstrated that aerobic glycolysis induced by metformin can significantly promote angiogenesis and neurogenesis in the aged brain.

## Materials and Methods

### Experimental animals

C57BL/6 SPF mice (7–8 months or 8 weeks of age) were obtained from Beijing Vital River Laboratory Animal Technology (Beijing, China). All mice were housed five per cage, maintained on a 12-h light/dark schedule, and allowed free access to food and water. Animal protocols were approved by the Animal Research Committee of Tongji University School of Medicine (Shanghai, China). Experimental mice were treated with 10 mg/kg metformin by tail vein injection or oral administration (higher concentration metformin was 25 mg/kg, lower concentration metformin was 1 mg/kg) or VEGF (80 ng/kg) recombinant protein by tail vein injection every two days.

### Morris water maze task

The MWM test was used to measure hippocampal-based spatial memory and learning functions. The MWM apparatus consisted of a circular pool (1.2-m diameter) that contained water maintained at 24–26°C. A clear, circular escape platform (11-cm diameter) was submerged approximately 1.5 cm below the water surface. To escape from the water, mice had to find the hidden escape platform. Each acquisition trial was started by placing the mouse in the water facing the wall of the tank. The training protocol consisted of 5 days (four trials per day). For each trial, the animal was placed into the maze near one of four possible points: north, south, southeast, or northwest. The location was determined randomly for each trial. During each trial, the animal was given 60 s to locate the submerged platform. If a mouse did not locate the platform, it was gently led to the platform. After either finding or being led to the platform, the animal was left on the platform for 20 s to get familiarized with its location with respect to visual cues. Animals were tested in squads of six to eight mice, with all treatment groups represented within each testing squad. The inter-trial interval for each mouse was approximately 20 min. The mice were subjected to probe trials 24 hours after training. During probe trials, the platform was removed and mice were allowed to swim in the pool for 60 s. A camera mounted on the ceiling in the center of the pool was used to track the swim route of the mouse. Data were collected using a computerized animal tracking system (EthoVision XT Base; Noldus 2020 Software, Nottingham, UK), which recorded the path length, swim speed, and time spent in each quadrant of the pool, as well as the time taken to reach the platform (latency). The time spent in each quadrant was recorded. Trials were consistently performed between 1 pm and 5 pm.

### Cerebral blood flow monitored by Moor laser Doppler imaging scanner

CBF was monitored at the site of the cranial window. Moor laser Doppler imaging (LDI) with a Moor LDI scanner (Moor Instruments, Wilmington, DE, USA) was performed with the camera in the center aiming at the animal’s brain at a height of 20 cm from the skull. Image acquisition size was 2.4 × 1.7 cm at a resolution of 172 × 118 pixels. Laser ramp speed was 4 ms/pixel. To achieve the highest CBF response, the right whiskers were stimulated for 1 min at 10 Hz. All mice were imaged in a separated scan of the same resolution, ramp speed, and image size. Increased CBF values relative to the resting level were expressed as fold-change relative to the control group.

### Cardiac perfusion of mice and tissue harvesting

Mice were deeply anaesthetized using tribromoethanol (Sigma, St. Louis, MO, USA). Blood was taken by removing the eyeball, and then centrifuged at 1000 ×*g*/min for 10 min at 4°C to obtain plasma for measuring fructosamine with fructosamine reagents (IDEXX Catalyst™ Test; Westbrook, ME, USA) on IDEXX VetTest equipment. Mice were cardiac perfused with 0.01 M phosphate-buffered saline (PBS) (Sigma). Brain hemispheres were placed on an ice-cold glass dissection plate and orientated in a sagittal plane. Right hemispheres were immersed in 4% PFA for cryostat sectioning and immunohistochemistry, and stored at −80°C until required.

### Cell culture

All protocols were approved by the Ethics Research Committee at Tongji University School of Medicine. NSCs were microdissected from the lateral ventricle walls (ependymal zone together with SEZ/SVZ) covering the striatum of mice, and cultured in DMEM/F12 supplemented with 2% B-27, to which 0.1% bFGF and human epidermal growth factor (hEGF) were added daily, at 37°C in a 5% CO2 environment.

### Immunostaining

Brain tissue slices (10-μm) or cultured cell samples were treated with 4% PFA for 20 min and stored at room temperature. After three washes with PBS, all slides were incubated in blocking buffer (3% bovine serum albumin, 5% normal donkey serum, and 0.3% Triton X-100 in PBS) for 1 h at room temperature. Slides or coverslips were immersed in primary antibody buffer (200 μL per slide or 40 μL per coverslip) for incubation overnight at 4°C. Primary antibodies used in this study include: anti-GFAP (1:1000; Dako, Glostrup, Denmark), anti-Tuj1 (1:500; R&D Systems, Minneapolis, MN, USA), anti-DCX (1:400; Cell Signaling Technology, Danvers, MA, USA), anti-BrdU (1:200; Abcam, Cambridge, UK), anti-Sox2 (1:200, Abcam), and anti-NeuN (1:300; Abcam). The following day, coverslips and slides were washed three times with PBS, and then incubated with appropriate Alexa488-, Alexa568-or Cy3-conjugated secondary antibodies (1:1000; Thermo Fisher Scientific, Waltham, MA, USA) for 1 h at room temperature, or lectin-FITC (Dylight, DL-1174, Vectorlabs, CA, USA) for 2 h. DAPI staining for 15 min was used to label nuclei.

### Spontaneous differentiation of NSCs *in vitro*

To examine spontaneous differentiation of NSCs microdissected from lateral ventricle walls (ependymal zone together with SEZ/SVZ) covering the striatum of mice treated with metformin, single-cell suspensions of NSCs were plated onto coverslips coated with polyornithine (0.25 mg/mL in PBS) and laminin (5 μg/mL in DMEM-F12) at a density of 2 × 10^5^ cells/cm^2^. When NSCs reached approximately 80% confluence, bFGF and hEGF were withdrawn, and cells were treated with 1 mmol/L metformin for 7 continuous days. Subsequently, cells were fixed with 4% PFA and immunostained with GFAP and Tuj1 antibodies.

### RNA sequencing and data analysis

Total RNA was extracted from blood by TRIzol (Invitrogen, Carlsbad, CA, USA) and quantified by a spectrophotometer (NanoDrop 1000; Thermo Fisher Scientific). Only samples that met our quality criteria (260/280 nm > 1.8) were included in experiments. All data analyses were based on clean data with high quality. Index of the reference genome was built using Bowtie v2.0.6 (bowtie-bio.sourceforge.net/bowtie2) and paired-end clean reads were aligned to the reference genome using TopHat v2.0.9 (ccb.jhu.edu/software/tophat). Reads per kilobase of transcript, per million mapped reads (RPKM) of each gene were calculated based on gene length and read counts mapped to the gene, and considering the effect of sequencing depth and gene length for read counts at the same time. All differentially expressed genes were used for heatmap analysis and Kyoto Encyclopedia of Genes and Genomes (KEGG; https://www.genome.jp/kegg) analysis. For KEGG analysis, a q-value < 0.05 was used as the threshold to determine significant enrichment of gene sets. RNA-Seq data were deposited at GSE140716.

### qRT-PCR

Total cellular RNA was isolated using TRIzol reagent (Invitrogen) according to the manufacturer’s instructions. RNA was further purified by DNAase treatment and removal kit (Ambion, Austin, TX, USA). RNA samples (1 μg each) were then reverse-transcribed into cDNA using a SuperScript III First-Strand Synthesis kit (Invitrogen) according to the manufacturer’s instructions. Real-time PCR was performed on a 7500 or Q7 real-time PCR system (Applied Biosystems, Foster City, CA, USA) using SYBR Premix Ex Taq with ROX (Bio-Rad, Hercules, CA, USA). Relative gene expression levels were normalized to β-actin and calculated as 2^−ΔΔCT^. The sequences of the primers used are listed in Table 1.

**Table 1.**
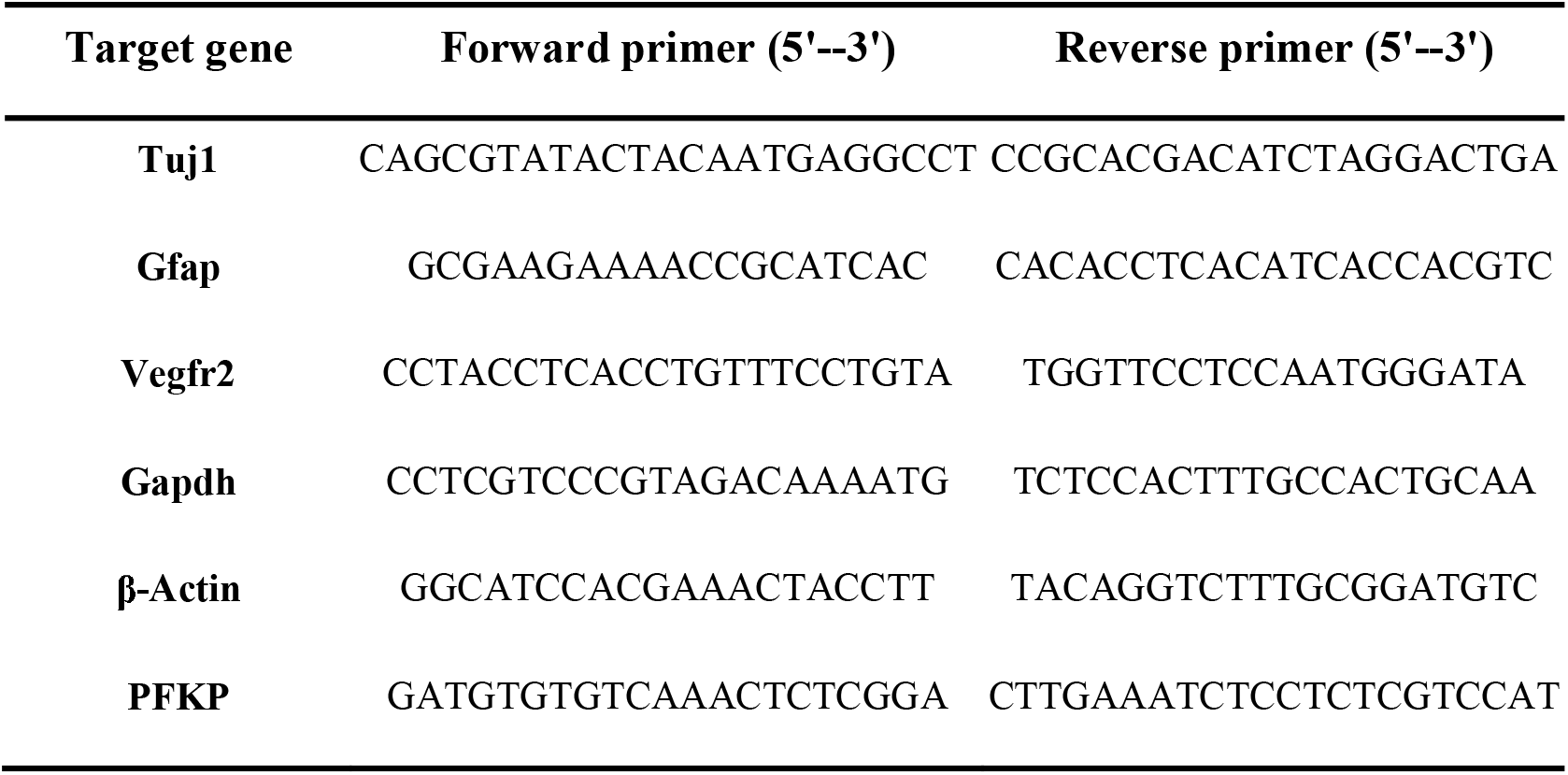
Primer sequences used in qRT-PCR.

### Quantification of immunohistochemical staining

Each experimental group contained at least three mice. Twelve serial sections (sagittal, 10 μm) per mouse were chosen for subsequent immunostaining, according to similar anatomical locations for each mouse.

### Human blood samples

Human blood samples were obtained from healthy volunteers. All subjects signed an informed consent form. The operation protocols were approved by the Ethics Research Committees from Huadong Hospital Affiliated to Fudan University and Tongji Hospital Affiliated to Tongji University.

### Graphics

Unless otherwise specified, plots were generated using R (https://www.R-project.org/) or GraphPad Prism 5 (GraphPad Software, San Diego, CA).

## Abbreviations

NSC: neural stem cell
GAPDH: glyceraldehyde-3-phosphate dehydrogenase
ROS: reactive oxygen species
SEZ/SVZ: subependymal zone/subventricular zone
MWM: Morris water maze
CBF: cerebral blood flow
Sox2: SRY-box 2
DCX: doublecortin
BrdU: bromodeoxyuridine
bFGF: basic fibroblast growth factor
PFA: paraformaldehyde
GFAP: glial fibrillary acidic protein
WGCNA: weighted gene co-expression network analysis
FOXO: Forkhead box
AMPK: AMP-activated protein kinase
qRT-PCR: quantitative real-time polymerase chain reaction
PFKP: phosphofructokinase
IA: iodoacetic acid
VEGFR2: vascular endothelial growth factor receptor 2.

## Author Contributions

L.H.L designed the study; Z.X.Q., S.J.Y and C.H performed and drafted the experiment; F.S.Y performed RNA-seq data analyses; L.H.L. revised the manuscript; L.H.L., L.Z.M., S.E.Y supervised the entire project; and all authors approved the final version of the manuscript.

## Conflicts of interest

All authors declare that no conflict of interest exists

## Funding

This work was supported by grants from the National Key Research and Development Program (2019YFC200187), the National Natural Science Foundation of China (Grant No 31671539, 31370214), and Major Program of Development Fund for Shanghai Zhangjiang National Innovtaion Demonstration Zone<Stem Cell Strategic Biobank and Stem Cell Clinical Technology Transformation Platform> (ZJ2018-ZD-004).

---

**Supplementary Figure 1. Metformin did not improve spatial learning and memory of 4-month-old mice in the Morris water maze test.** (A) Mean escape latency during 5 training days was not different in 4-month-old mice treated with metformin (*n* = 5) compared with the control group (*n* = 6). (B) Probe tests conducted 24 h after the acquisition phase indicated that the first time-to-platform was no different in 4-month-old mice treated with metformin than the control group. (C) Mean speed was also not different between the two groups. Ctrl: Control; Met: Metformin. The overall significance between two groups was determined by Student’s *t*-test. * *p* < 0.05, ** *p* < 0.01, *** *p* < 0.001, ns, not significant.

**Supplementary Figure 2. Concentration of plasma fructosamine was decreased in 8-month-old mice treated with metformin (*n* = 4) compared with controls (*n* = 3)**. Ctrl: Control; Met: Metformin; FRU: Fructosamine. The overall significance between two groups was determined by Student’s *t*-test. * *p* < 0.05, ** *p* < 0.01.

**Supplementary Figure 3. GAPDH levels positively correlate with cognitive levels in humans.** (A) Relative GAPDH mRNA expression in blood of young and aged people. (B) Relative GAPDH mRNA expression in blood sample from people of various ages. (C) Relative GAPDH mRNA expression in blood sample of old people (chronicle age > 70s) positively correlated with cognitive levels measured by the Mini-Mental State Examination. * *p* < 0.05, ** *p* < 0.01, *** *p* < 0.001, ns, not significant.

## References

1. Tarantini S, Tran CHT, Gordon GR, Ungvari Z, Csiszar A. Impaired neurovascular coupling in aging and Alzheimer’s disease: Contribution of astrocyte dysfunction and endothelial impairment to cognitive decline. Exp Gerontol. 2017;94:52–58. https://doi.org/10.1016/j.exger.2016.11.004 PMID: 27845201

2. Farzaneh A. Sorond SH, David H. Salat, Douglas N. Greve, Naomi D.L. Fisher. Neurovascular coupling, cerebral white matter integrity, and response to cocoa in older people. Neurology. 2013;81:904–909. https://doi.org/10.1212/WNL.0b013e3182a351aa PMID: 23925758

3. Matilde Balbi MG, Thomas A Longden, Max Jativa Vega, Benno Gesierich, Farida Hellal, Athanasios Lourbopoulos, Mark T Nelson and Nikolaus Plesnila. Dysfunction of mouse cerebral arteries during early aging. Journal of Cerebral Blood Flow & Metabolism. 2015:1–9. https://doi.org/10.1038/jcbfm.2015.107 PMID: 26058694

4. Farzaneh A Sorond, Dan K Kiely, Andrew Galica, Nicola Moscufo, Jorge M Serrador, Ike Iloputaife, Svetlana Egorova, Elisa Dell’Oglio, Dominik S Meier, Elizabeth Newton, William P Milberg, Charles R G Guttmann, Lewis A Lipsitz. Neurovascular coupling is impaired in slow walkers: The MOBILIZE Boston Study. Annals of Neurology. 2011;70(2):213–220. https://doi.org/10.1002/ana.22433 PMID: 21674588

5. Eelen G, de Zeeuw P, Simons M, Carmeliet P. Endothelial Cell Metabolism in Normal and Diseased Vasculature. Circulation Research. 2015;116(7):1231–1244. https://doi.org/10.1161/circresaha.116.302855 PMID: 25814684

6. Masoud Tavazoie, Lieven Van der Veken, Violeta Silva-Vargas, Marjorie Louissaint, Lucrezia Colonna, Bushra Zaidi, Jose Manuel Garcia-Verdugo, Fiona Doetsch. A specialized vascular niche for adult neural stem cells. Cell stem cell. 2008;3(3):279–288. https://doi.org/10.1016/j.stem.2008.07.025 PMID: 18786415

7. Cécile Batandier, Bruno Guigas, Dominique Detaille, M-Yehia El-Mir, Eric Fontaine, M Rigoulet, Xavier M Leverve. The ROS Production Induced by a Reverse-Electron Flux at Respiratory-Chain Complex 1 is Hampered by Metformin. Journal of Bioenergetics and Biomembranes. 2006;38(1):33–42. https://doi.org/10.1007/s10863-006-9003-8 PMID: 16732470

8. Westhaus A, Blumrich EM, Dringen R. The Antidiabetic Drug Metformin Stimulates Glycolytic Lactate Production in Cultured Primary Rat Astrocytes. Neurochem Res. 2017;42(1):294–305. https://doi.org/10.1007/s11064-015-1733-8 PMID: 26433380

9. Björn Neumann, Roey Baror, Chao Zhao, Michael Segel, Sabine Dietmann, Khalil S Rawji, Sarah Foerster, Crystal R McClain, Kevin Chalut, Peter van Wijngaarden, Robin J M Franklin. Metformin Restores CNS Remyelination Capacity by Rejuvenating Aged Stem Cells. Cell stem cell. 2019;25(4):473–85 e8. https://doi.org/10.1016/j.stem.2019.08.015 PMID: 31585093

10. Jing Wang, Denis Gallagher, Loren M DeVito, Gonzalo I Cancino, David Tsui, Ling He, Gordon M Keller, Paul W Frankland, David R Kaplan, Freda D Miller. Metformin activates an atypical PKC-CBP pathway to promote neurogenesis and enhance spatial memory formation. Cell stem cell. 2012;11(1):23–35. https://doi.org/10.1016/j.stem.2012.03.016 PMID: 22770240

11. Katsimpardi L, Litterman NK, Schein PA, Miller CM, Loffredo FS, Wojtkiewicz GR, et al. Vascular and neurogenic rejuvenation of the aging mouse brain by young systemic factors. Science. 2014;344(6184):630–634. https://doi.org/10.1126/science.1251141 PMID: 24797482

12. Tavazoie M, Van der Veken L, Silva-Vargas V, Louissaint M, Colonna L, Zaidi B, John W Chen, Richard T Lee, Amy J Wagers, Lee L Rubin. A Specialized Vascular Niche for Adult Neural Stem Cells. Cell Stem Cell. 2008;3(3):279–288. https://doi.org/10.1016/j.stem.2008.07.025 PMID: 24797482

13. Sirover MA. New nuclear functions of the glycolytic protein, glyceraldehyde-3-phosphate dehydrogenase, in mammalian cells. J Cell Biochem. 2005;95(1):45–52. https://doi.org/10.1002/jcb.20399 PMID: 15770658

14. Sen N, Hara MR, Kornberg MD, Cascio MB, Bae BI, Shahani N, Bobby Thomas, Ted M Dawson, Valina L Dawson, Solomon H Snyder, Akira Sawa. Nitric oxide-induced nuclear GAPDH activates p300/CBP and mediates apoptosis. Nat Cell Biol. 2008;10(7):866–873. https://doi.org/10.1038/ncb1747 PMID: 18552833

15. Chang C, Su H, Zhang D, Wang Y, Shen Q, Liu B, Rui Huang, Tianhua Zhou, Chao Peng, Catherine C L Wong, Han-Ming Shen, Jennifer Lippincott-Schwartz, Wei Liu. AMPK-Dependent Phosphorylation of GAPDH Triggers Sirt1 Activation and Is Necessary for Autophagy upon Glucose Starvation. Mol Cell. 2015;60(6):930–940. https://doi.org/10.1016/j.molcel.2015.10.037 PMID: 26626483

16. Zheng L, Roeder RG, Luo Y. S Phase Activation of the Histone H2B Promoter by OCA-S, a Coactivator Complex that Contains GAPDH as a Key Component. Cell. 2003;114(2):255–266. https://doi.org/10.1016/s0092-8674(03)00552-x PMID: 12887926

17. Azam S, Jouvet N, Jilani A, Vongsamphanh R, Yang X, Yang S, Ramotar D. Human glyceraldehyde-3-phosphate dehydrogenase plays a direct role in reactivating oxidized forms of the DNA repair enzyme APE1. J Biol Chem. 2008;283(45):30632–30641. https://doi.org/10.1074/jbc.M801401200 PMID: 18776186

18. Alexander A Shestov XL, Zheng Ser, Ahmad A Cluntun, Yin P Hung, Lei Huang, Dongsung Kim, Anne Le, Gary Yellen, John G Albeck, Jason W Locasale. Quantitative determinants of aerobic glycolysis identify flux through the enzyme GAPDH as a limiting step. eLife. 2014;3:1–18. https://doi.org/10.7554/eLife.03342 PMID: 25009227

19. Folstein. MF FS, McHugh. PR. “Mini-mental state”: A practical method for grading the cognitive state of patients for the clinician. J Psychiatr Res. 1975;12:189–198. https://doi.org/10.1016/0022-3956(75)90026-6 PMID: 1202204

20. Duarte JMN, Do KQ, Gruetter R. Longitudinal neurochemical modifications in the aging mouse brain measured in vivo by 1H magnetic resonance spectroscopy. Neurobiology of Aging. 2014;35(7):1660–1668. https://doi.org/10.1016/j.neurobiolaging.2014.01.135 PMID: 24560998

21. Ennaceur A, Michalikova S, van Rensburg R, Chazot PL. Detailed analysis of the behavior and memory performance of middle-aged male and female CD-1 mice in a 3D maze. Behav Brain Res. 2008;187(2):312–326. https://doi.org/10.1016/j.bbr.2007.09.025 PMID: 17983672

22. Francia N, Cirulli F, Chiarotti F, Antonelli A, Aloe L, Alleva E. Spatial memory deficits in middle-aged mice correlate with lower exploratory activity and a subordinate status: role of hippocampal neurotrophins. Eur J Neurosci. 2006;23(3):711–728. https://doi.org/10.1111/j.1460-9568.2006.04585.x PMID: 16487153

23. Thangthaeng N, Rutledge M, Wong JM, Vann PH, Forster MJ, Sumien N. Metformin Impairs Spatial Memory and Visual Acuity in Old Male Mice. Aging Dis. 2017;8(1):17–30. https://doi.org/10.14336/AD.2016.1010 PMID: 28203479

24. de Jager J, Kooy A, Lehert P, Wulffele MG, van der Kolk J, Bets D, Joop Verburg, Ab J M Donker, Coen D A Stehouwer. Long term treatment with metformin in patients with type 2 diabetes and risk of vitamin B-12 deficiency: randomised placebo controlled trial. Bmj. 2010;340:c2181. https://doi.org/10.1136/bmj.c2181 PMID: 20488910

25. Martin-Montalvo A, Mercken EM, Mitchell SJ, Palacios HH, Mote PL, Scheibye-Knudsen M, Gomes PA, Ward MT, Minor KR, Blouin MJ, Schwab M, Pollak M, Zhang Y, YuY, Becker KG, Bohr VA, Ingram DK, Sinclair DA, Wolf NS, Spindler SR, Bernier M, Cabo R. Metformin improves healthspan and lifespan in mice. Nature Communications. 2013;4(1). https://doi.org/10.1038/ncomms3192 PMID: 23900241

26. Rebecca M. Ruddy KVA, Cindi M. Morshead. Age- and sex-dependent effects of metformin on neural precursor cells and cognitive recovery in a model of neonatal stroke. Sci Adv. 2019;5 (eaax1912):1–10. https://doi.org/10.1126/sciadv.aax1912 PMID: 31535024

27. Bensalem J, Servant L, Alfos S, Gaudout D, Layé S, Lafenetre P. Dietary Polyphenol Supplementation Prevents Alterations of Spatial Navigation in Middle-Aged Mice. Frontiers in Behavioral Neuroscience. 2016;10,9. https://doi.org/10.3389/fnbeh.2016.00009 PMID: 26903826

28. Newman LA, Korol DL, Gold PE. Lactate produced by glycogenolysis in astrocytes regulates memory processing. PLoS One. 2011;6(12):e28427. https://doi.org/10.1371/journal.pone.0028427 PMID: 22180782

29. Suzuki A, Stern SA, Bozdagi O, Huntley GW, Walker RH, Magistretti PJ, Alberini CM. Astrocyte-neuron lactate transport is required for long-term memory formation. Cell. 2011;144(5):810–823. https://doi.org/10.1016/j.cell.2011.02.018 PMID: 21376239

30. Yu P, Wilhelm K, Dubrac A, Tung JK, Alves TC, Fang JS, Xie Y, Zhu J, Chen Z, Smet FD, Zhang J, Jin SW, Sun L, Sun H, Kibbey RG, Hirschi KK, Hay N, Carmeliet P, Chittenden TW, nne Eichmann A, Potente M, Simons M. FGF-dependent metabolic control of vascular development. Nature. 2017;545(7653):224–228. https://doi.org/10.1038/nature22322 PMID: 28467822

31. Qin Shen SKG, Li Jin, Nithin Karanth, Yu Sun, Natalia Abramova, Peter Vincent, Kevin Pumiglia, Sally Temple. Endothelial Cells Stimulate Self-Renewal and Expand Neurogenesis of Neural Stem Cells. Science. 2004;304(5675): 1338–1340. https://doi.org/10.1126/science.1095505 PMID: 15060285

32. S. Miyama TT, R.S. Nowakowski and V.S. Caviness, Jr. A Gradient in the Duration of the G1 Phase in the Murine Neocortical Proliferative Epithelium. Cerebral Cortex. 1997;7(7):678–689. https://doi.org/10.1093/cercor/7.7.678 PMID: 9373022

