## Supplementary figures and images for "Metformin improves cognition of aged mice by promoting cerebral angiogenesis and neurogenesis"

### Supplementary Figure.1

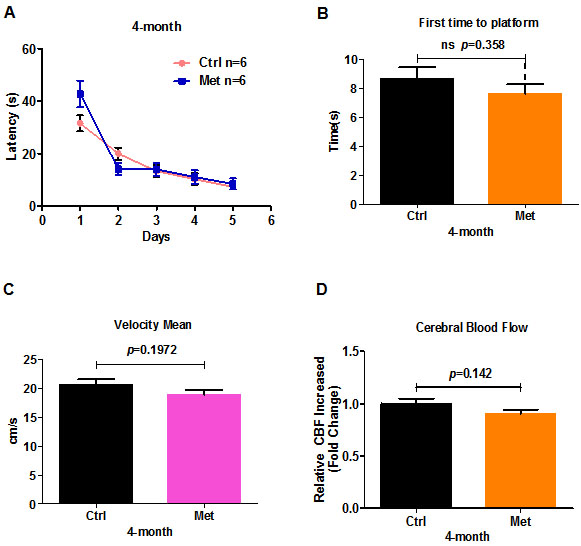

### Supplementary Figure.2

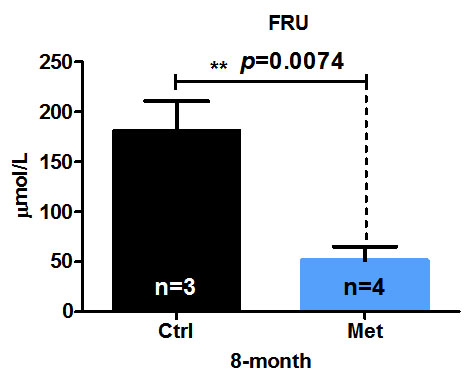

### Supplementary Figure.3

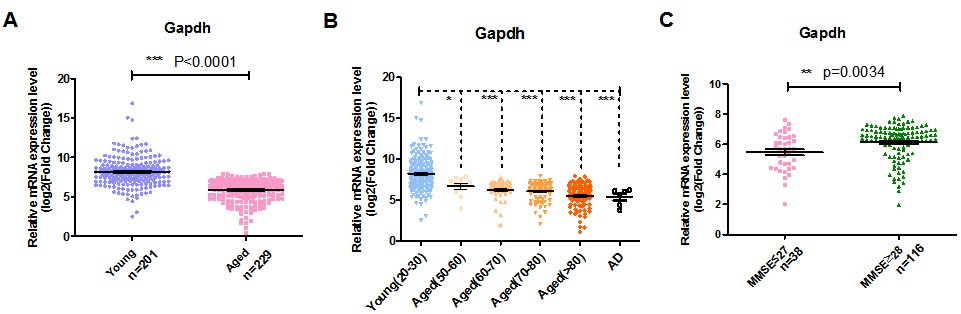
